# Cascaded Antithetic Integral Feedback for Enhanced Stability and Performance

**DOI:** 10.1101/2024.07.31.605983

**Authors:** Armin M. Zand, Ankit Gupta, Mustafa Khammash

## Abstract

Precise intracellular regulation and robust perfect adaptation can be achieved using biomolecular integral controllers and it holds enormous potential for synthetic biology applications. In this letter, we consider the cascaded implementation of a class of such integrator motifs. Our cascaded integrators underpin proportional-integral-derivative (PID) control structures, which we leverage to suggest ways to improve dynamic performance. Moreover, we demonstrate how our cascaded strategy can be harnessed to enhance robust stability in a class of uncertain reaction networks. We also discuss the genetic implementation of our controllers and the natural occurrence of their cascaded sequestration pairs in bacterial pathogens.

## I. Introduction

THE phenomenon of robust perfect adaptation (RPA), a hallmark of essential natural processes from the cellular to the organ level, underpins many vital life systems [1]– [3]. Achieving it through genetic devices promises advancements in synthetic biology applications (see [2] and references therein). Model-based designs of static feedback controllers, particularly integral feedback control (IFC), realizable through chemical reaction networks (CRNs), are promising initial steps. Recently, there has been significant interest in the characterization, design, and instantiation of biomolecular controllers and feedback systems for robust gene-expression control [4]–[15]. Antithetic integrators [7], in particular, have successfully achieved synthetic RPA in both bacteria [6], [9] and mammalian cells [15], [16].

Since its introduction in [7], the antithetic integral feedback controller (AIC) and its variants have received significant attention and have been studied theoretically from both deterministic and stochastic perspectives [17]–[28]. At its core, this motif uses sequestration between two species to compute the integral of the tracking error. Under standard assumptions such as set-point admissibility and stability, this *core* antithetic controller tightly regulates the average concentration of a species of interest across a cell population [7].

It goes without saying that proportional-integral-derivative (PID) control has been the backbone of industrial control technology for its practical advantages [29], [30]. Using the core AIC, a rich repertoire of variant motifs have been proposed in the literature that encode PID control structures to enhance the dynamic performance [17]–[ Simulations suggest their effectiveness in tuning nominal transient characteristics. Gain inseparability is a known issue in nonlinear PID realizations through CRNs [17], [19], inherent to local analysis, where PID gains depend on both the design parameters and steady-state values. This complicates fair comparisons of different PID motifs, as their flexibility in gain adjustment through biomolecular parameter fine-tuning remains process- and disturbance-specific. Thus, having different topologies that expand the design space, offer practical advantages, or provide greater tuning flexibility is both useful and crucial. Recent works have also studied the stability of antithetic control in certain plants [22]–[27] Suggested stabilizing approaches hitherto include exploiting the controller constant degradation/dilution, molecular buffering, low-gain actuation, and the rein mechanism.

Building upon the core AIC motif [7], this letter introduces a higher-dimensional implementation of it—a controller motif we call “cascaded” AIC. In Section II, we first present a generalized formulation of this cascaded motif, followed by a representative minimal controller. The minimal cascaded controller motif remains the main focus of our study. We also discuss how the cascaded sequestration reactions, necessary components for our designs, naturally occur in certain bacteria. The contributions of this paper are to explore our proposed biocontrollers and their control theoretic features. Previous works [19], [21] have shown that the core AIC follows a filtered PI (resp. I) control structure when actuating via the sensor (resp. reference) controller species. In comparison, our cascaded structure encodes a filtered PID (resp. PI) mechanism when the actuation arrangement is the same. These underlying filtered PI and PID control structures are, therefore, intrinsic to our controller architecture itself. In Section III, we demonstrate how exploiting this novel architecture offers practical means and flexibility to improve nominal control performance.

Cellular systems are known for their inherent noise and complexity [31]. Considering large parametric/structural uncertainties in their nominal deterministic models is thus relevant, if not imperative. Ideally, the closed-loop system should be insensitive to these uncertainties. The risk of instability, however, is a known drawback of feedback systems [29], exacerbated when large uncertainties are present. Section IV delves into the analysis of stability robustness, characterizing a wide class of uncertain reaction networks that can be stabilized by our cascaded controller. Our approach leverages the tuning of certain constant inflow (zeroth-order) rates to lower the integrator gain. Remarkably, this method offers a flexible tuning knob for achieving robust stability in a biologically convenient manner, potentially minimizing the need for complex protein engineering that might otherwise be required if, e.g., design parameters associated to first or second-order rates were to be tuned. Before concluding this letter, we also discuss potential biomolecular realizations of our introduced controllers.

## II. Cascaded Antithetic Integrators

### A. Generalized Cascaded AIC Motifs

Let us assume that an uncertain, poorly-known process network of interest, called 𝒳, is given. The control objective is a regulation task: designing a controller that steers the concentration of a target species within 𝒳, say **X**_**n**_, to a pre-defined set-point, provided proper sensing and actuation. This controller acts on a known input species, **X**_**1**_. The regulation must be robust to process variations, and the set-point value should be inducible by the controller. Capable of achieving such regulation (RPA), our proposed controller, named cascaded AIC, is schematically illustrated in Fig. 1. For the *q*thorder cascaded AIC, we cascade *q* − 1 different annihilation modules through *q* controller species, **Z**_**1**_ to **Z**_**q**_. Following the generic reaction 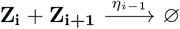 for *i* ∈ {1, · · ·, *q* − 1}, each controller species **Z**_**i**_ forms a sequestration pair with **Z**_**i+1**_, undergoing co-degradation at a rate *η*_*i*−1_. The sensor species **Z**_**2**_ measures the current value of **X**_**n**_, which is used for computing the integral error and offsetting it. Except for **Z**_**2**_ and **Z**_**1**_, every controller species **Z**_**i**_ undergoes constant production at a rate *µ*_*i*−2_, as shown in Fig. 1A.

**Fig. 1.**
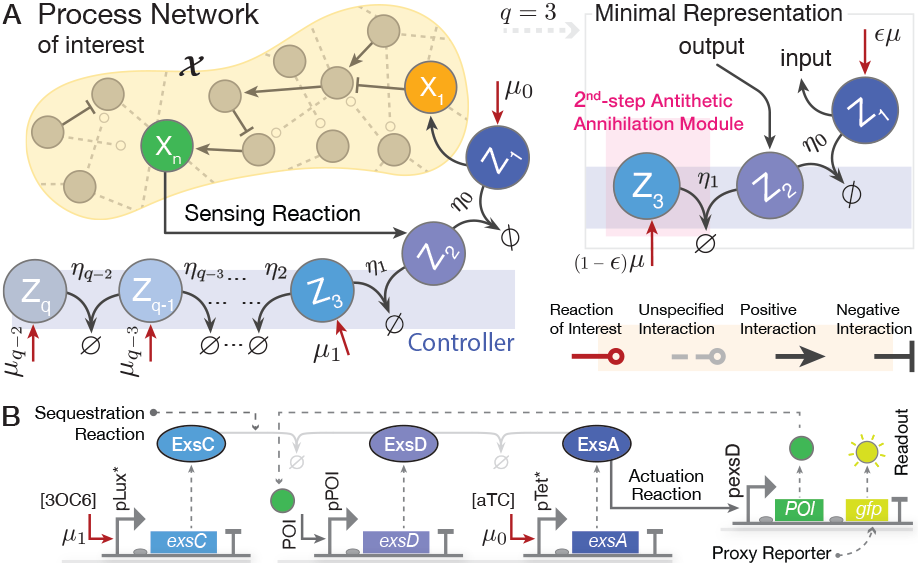
Cascaded antithetic integral feedback control topology. (A) The general depiction of the motif is on the left panel, with the minimal representation featuring three active controller species on the right. The controller closes the loop by sensing from and actuating on an uncertain, poorly-known process network. (B) An example circuit to realize the minimal cascaded AIC motif, based on the regulatory genes involved in the T3SS of *P. aeruginosa* bacteria [32]. The controller is employed to regulate a one-species process comprising the protein of interest (POI). The POI acts as an activator for **Z**_**2**_. It is inserted in an operon including a fluorescent protein as a proxy for readout, whose expression level is proportional to that of the POI. pLux* and pTet* control the expressions of **Z**_**3**_ **an**d **Z**_**1**_, respectively. The inducers 3OC6 and aTC are used to tune the activity of these two promoters. By adjusting their availability in the culture medium, one can tune both inflow rates *µ*_0_ and *µ*_1_. The mapping between the biological species and Z_1_ to Z_3_ is indicated by matching colors.

We do not specify functional roles for the resulting sequestration complexes, which may be active or inactive depending on the context. Nonetheless, their inclusion would not change the overall capacity of our controller to achieve RPA. By defining the integral variable 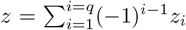,it is seen that 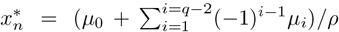,Throughout, we use lowercase letters to represent the concentrations of the corresponding species denoted by bold, capital letters, and superscript asterisks to denote steady-state values. A closedloop dynamical model for this generalized cascaded antithetic controller is reported in Supplementary Material, Section S1. A list of necessary conditions for set-point admissibility is included therein, which are satisfied if the constitutive production rate of each **Z**_**i**_ is higher than that of its adjacent **Z**_**i+1**_ for *i* ≥ 3, meaning that for the case *q* = 3, the controller dynamics impose no constraint. In providing our dynamic models, we assume deterministic models, the law of mass action, and negligible effects of constant controller degradation/dilution.

### B. The Minimal Representation and Its Dynamical Model

The minimal cascaded AIC we obtain from the general motif by setting *q* = 3. Depicted in Fig. 1A, this motif consists of three species and two intertwined annihilation modules. The augmentation of the additional annihilation module changes the control structure. Later sections will discuss how this enables robustness and performance improvement. The resulting closed-loop dynamical behavior can be described by

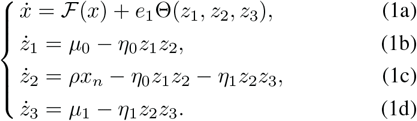

Recent preprint [28] explores noise reduction with a similar controller motif for a two-species process, using different actuation and sensing mechanisms and stochastic simulations. In contrast, our analysis focuses on the control theoretic properties, where we present new results regarding stability and performance in a generalized deterministic framework. Compared to the core AIC [7], a new species, **Z**_**3**_, is added to the controller side, forming a sequestration pair with **Z**_**2**_. The dynamical behavior of the open-loop circuit 𝒳, which comprises *n* ≥ 1 distinct species **X**_**1**_ to **X**_**n**_, is described by the smooth vector field ℱ(*x*) := [*f*_1_, …, *f*_*n*_]^*T*^ : ℝ^*n*^ → ℝ^*n*^. Within the controller network, two separate annihilation reactions occur with potentially different propensities, *η*_0_ and *η*_1_. The function Θ represents the cumulative effect of the control input applied to X_1_. For now, we allow the controller species **Z**_**1**_ to **Z**_**3**_ to freely act on the process input X_1_, though Θ may need to conform to certain specifications in later sections. By introducing the positive constants ϵ *<* 1 and *µ*, we parametrize the inflow rates *µ*_*i*_ as follows throughout this letter

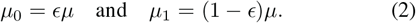

In this case, it follows that 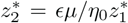 and 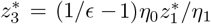. Setting ϵ = 1 and *z*_3_(0) = 0 neutralizes the extra annihilaiton module, reducing the controller to the core AIC. We use this fact when comparing our designs with the core AIC. Under the assumptions of set-point admissibility and closed-loop stability, the output of interest regulates to the value *µ/ρ* at steady state. The former assumption, which corresponds to the feasibility of having a positive equilibrium point reachable from its neighborhood, remains an assumption throughout this letter. However, Section IV will provide guarantees of closed-loop stability for a class of process networks.

### C. Cascades of Sequestration Naturally Appearing in Bacteria

We observe that the sequestration pairs match up in cascaded forms among native proteins found in various bacterial pathogens [32], [33], mainly in their type-III secretion system (T3SS) that enables them to infect human cells. For example, in *P. aeruginosa* bacteria, we observe cascaded sequestrations up to the fourth order between the proteins ExsA, ExsD, ExsC, and ExsE. We have provided further examples in Supplementary Material, Table S1. The natural occurrence of the sequestration pairs required for our control topologies in bacterial genomes can simplify their incorporation. This emphasizes that higher-order complex control designs may not necessarily come with additional barriers to implementation.

## III. Exploiting the Underlying pi and pid Control Structures TO Improve NominaL Performance

In this section, we focus on the closed-loop dynamic performance and explore the added design flexibility offered by the minimal cascaded AIC motif. Specifically, we investigate whether the structural difference from the core AIC, brought about by the addition of the extra annihilation module, provides this flexibility. To do this, we analyze the controller transfer function standalone. We allow each Z_i_ to be involved in Θ, which expands our design flexibility. Recall that Θ(*t*) represents the applied actuation signal. Our small-signal analysis uncovers that, enabled by certain arrangements of Θ, the minimal cascaded AIC encodes PI and PID compensatory structures. The Laplace domain transformation of the control input Θ linearized about the fixed point follows

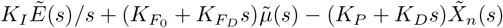

filtered in series through a second-order low-pass filter. Here, *K*_*P*_, *K*_*I*_, *K*_*D*_, and 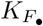 are the proportional, integral, derivative, and feedforward gains. 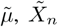, and 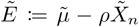,are the reference, output, and tracking error signals, respectively.

The above gains are provided in Supplementary Material, Section S2B in terms of biomolecular parameters. In particular, for *K*_*P*_, *K*_*I*_, and *K*_*D*_, we have the following expressions *K*_*P*_ = *ρ* (*σ*_11_*β*_0_*/α*_0_ − *σ*_21_(*α*_0_ + *β*_1_*/α*_1_) + *σ*_31_*α*_1_) */ω, K*_*I*_ = (*σ*_11_*β*_0_*β*_1_*/α*_0_*α*_1_ − *σ*_21_*α*_0_*β*_1_*/α*_1_ + *σ*_31_*α*_1_*α*_0_)*/ω*, and *K*_*D*_ = −*σ*_21_*ρ/ω*, respectively, where 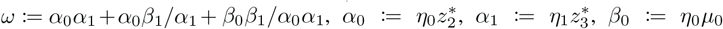, and *β*_1_ := *η*_1_*µ*_1_. The constants *σ*_*i*1_ are determined by the biomolecular reactions governing Θ, each representing the linearized effect of the actuation signal from **Z**_**i**_ to **X**_**1**_, that is, *σ*_*i*1_ := ∂Θ*/*∂*z*_*i*_ evaluated at the fixed point. For PID control with positive *K*_*D*_, these gains necessitate having a negative actuation from **Z**_**2**_. By rearranging the actuation terms in a way *σ*_11_ ≥ 0, *σ*_21_ ≤ 0, and *σ*_31_ ≥ 0, three different PI and four PID variant motifs can be obtained that neither require satisfying parametric conditions to maintain positive PID gains nor introduce additional nonminimum-phase zeros into the closed-loop system. See Supplementary Material, Section S2 for details and their depiction. Interestingly, with only a single **Z**_**i**_ actuating—specifically, a negative (resp. positive) actuation from **Z**_**2**_ (resp. **Z**_**1**_ or **Z**_**3**_)— one can realize a filtered PID (resp. PI) control structure. Considering the note in Section II-C, this can offer practical advantages when engineering as few reaction channels as possible between the process and controller is a design constraint.

Specific designs for Θ, e.g., those involving Hill kinetics, may offer greater flexibility in tuning the gains and cutoff frequencies, thereby enhancing the tunability of both disturbance rejection and tracking performances. Process variations, however, influence these gains, as mentioned in Section I. Despite this, we may still be able to exploit the extra degrees of freedom that the encoded PI and PID architectures offer, to significantly improve the transient dynamics of the *nominal* system. Fig. 2 confirms this for a particular Θ across three of the motifs. We observe that the core AIC response for the same process and set-point is unstable, whereas the control signals generated by its cascaded versions stabilize the system. The comparison of different designs presented in Fig. 2 show that non-zero values for *θ*_21_ and *θ*_31_, corresponding to the type-III PI and PID motifs, result in overall better nominal performance indices, demonstrating significant tunability over the transient characteristics. We present additional numerical simulations in Fig. S4 for a different function Θ involving the active degradation of **X**_**1**_ by **Z**_**2**_, which similarly suggest that cascaded motifs, particularly the PID designs, offer superior dynamic performance tunability compared to the core AIC.

**Fig. 2.**
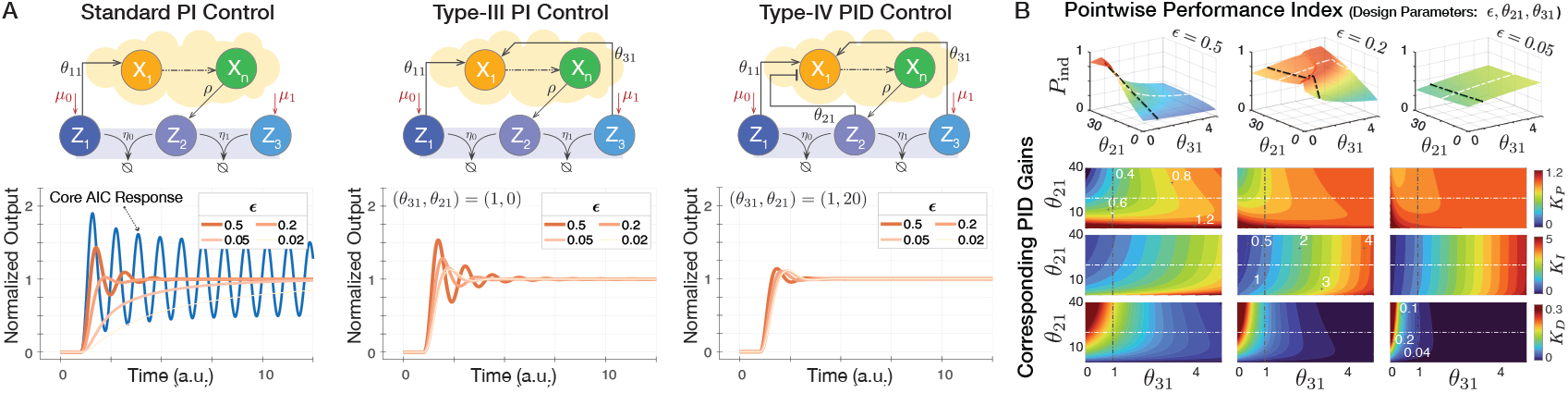
Exploiting the filtered PI and PID control structures brought about by the cascaded implementation of the antithetic IFC provides a practical means for improving nominal control performance. The linear model of our cascaded AIC follows a filtered PI (PID) control structure when actuating only by **Z**_**1**_ (**Z**_**2**_). Different actuation rearrangements can lead to higher degrees of tunability, resulting in different types of PI and PID mechanisms. The regulated process is a two-species gene expression example where 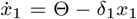 and 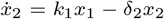 (A) For four different arrangements of the design parameter *ϵ*, we illustrate the performance improvement for two PI and one PID controller designs, comparing them with each other and the core AIC. The core antithetic response, which corresponds to *ϵ* = 1, shows instability. Here, Θ := *θ*_31_*z*_3_ + *θ*_11_*z*_1_*/*(1 + *θ*_21_*z*_2_), *θ*_11_ = 6, and the process parameters are fixed. (B) The pointwise performance index for variations in the design parameters *θ*_21_ and *θ*_31_ for three choices of *ϵ*. The 2D contour plots indicate the proportional (*K*_*P*_), integral (*K*_*I*_), and derivative (*K*_*D*_) gains corresponding to the design parameters. The regions within which we obtain *P*_ind_ are chosen such that they are locally stable. From the performance index plots for the above control circuits, one can pick an optimal pair for *θ*_21_ and *θ*_31_.

For our simulation case studies throughout this letter, we consider an *n*-species unimolecular activation cascade as the controlled process (see Fig. 3A), which are good models of known intracellular processes, e.g., gene expression (with maturation stages) activated through signaling pathways. We consider ℱ(*x*) = *Ax* where *A* := [*a*_*ij*_] ∈ ℝ ^*n*×*n*^ is a lower-triangular Metzler matrix with every element set to zero except *a*_*ii*_ = −*δ*_*i*_ and *a*_(*i*+1)*i*_ := *k*_*i*_. *δ*_*i*_ and *k*_*i*_ represent the process degradation and production rate constants. For the choices of Θ in our simulations, every positive set-point *µ/ρ* is admissible for this process model controlled by the controller given in (1). Numerical values and details regarding all figures are available in Supplementary Material, Section S3. The transient performance is indexed by a measure *P*_ind_, which is a weighted quantity of calculated overshoot (*Y*_ovr_), settling time (*T*_set_), and rise time (*T*_ris_) from time trajectory data. This measure calculates the following value: *P*_ind_ := 1*/*(1+*w*_1_*T*_set_ +*w*_2_*T*_ris_ + *w*_3_*Y*_ovr_). The best (worst) achievable *P*_ind_ is unity (zero). The weights are set to *w*_1_ = 0.1, *w*_2_ = 0.1, *w*_3_ = 10, prioritizing the damping of overshoot. Minimizing the response overshoot is particularly important in therapeutic applications, as it helps maintain drug exposure within the therapeutic window and avoid concentrations that could lead to toxicity.

**Fig. 3.**
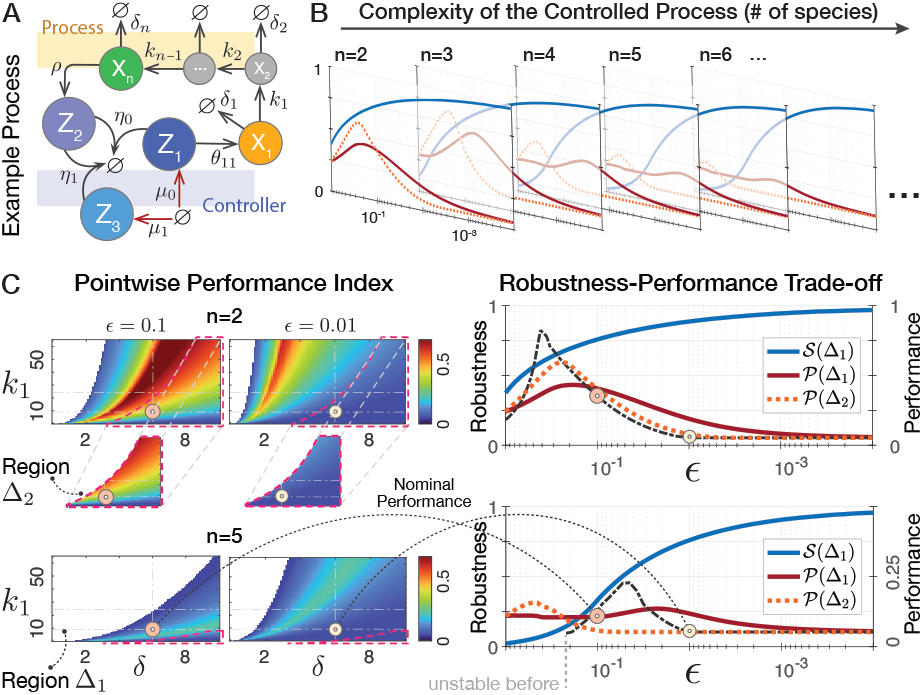
The trade-off between stability robustness to process variations and control performance is illustrated using two metrics, 𝒮(Δ) and 𝒫(Δ). The argument Δ identifies the uncertain parametric region for computation. Both metrics operate on the pointwise results of the stability status and performance index over Δ, while keeping the other parameters fixed at their nominal values. The metric 𝒮 ∈ [0, 1] computes the stable fraction within Δ, whereas 𝒫 ∈ [0, 1] averages the pointwise performance indices (*P*_ind_ defined in Fig. 2) over all stable points within Δ. The dash-dotted black lines indicate the nominal performance index. The region Δ_1_ represents the entire rectangle within which the process parameters *k*_1_ and *δ*_*i*_ =: *δ* vary. Δ_2_ is the common stable region across the considered variations of *ϵ*. It is a subset of Δ_1_ in which every point is locally stable given all uncertain regions of *k, δ*, and *ϵ*.

## IV. Robust StabilitY Analysis

The nominal performance improvement discussed in the previous section may come at the cost of compromised robust stability when uncertainties are taken into account. This can lead to trade-offs between stability robustness and transient performance [13], [29]. Such hard limits are often imposed by the introduction of feedback, particularly relevant in IFC systems due to Bode’s integral theorem [22], [29]. Simulations provided in Fig. 3 illustrate this trade-off for a set of nominal parameters and bounded uncertainties. As shown, tuning the design parameters—here *c*, which parameterizes the induced set-point distributed among Z_1_ and Z_3_ as in (2)—can improve both the performance and stability compared to the core AIC. However, for smaller *c*s the performance indicators reach their lowest points, while the closed-loop is stabilized across almost the entire uncertain range. Fig. 3B shows that adding more chains to the cascade 𝒳 compromises the average performance but shifts its peak to a higher stability score, indicating the role that the process complexity might play in this trade-off.

This section discusses tuning strategies to improve robust stability. We are mainly interested in achieving this through tuning *µ*_*i*_, which can significantly simplify design constraints. We carry a linear perturbation analysis local to an equilibrium point Σ and prove its stability for sufficiently small choices of *c*. Let us denote the proper transfer function associated with the open-loop circuit 𝒳 by *P* (*s*) := *N* (*s*)*/D*(*s*), with *N* and *D* being polynomials in *s* ∈ ℂ. It follows 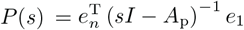,where *A*_p_ := ∂ℱ|_Σ_ is the Jacobian matrix at the steady-state Σ and *I* is the identity matrix.

### Assumptions.

*The following hold for the closed-loop (1):*

*(A1) The set-point* 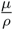*is admissible, implying the existence of an isolated fixed point* 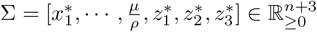.

*(A2) The function* Θ *is independent of z*_2_ *and z*_3_.

### Theorem 1.

*Suppose (A1)-(A2) are satisfied. Define σ*_11_ := ∂Θ*/*∂*z*_1_|_Σ_ *and N* ^t^(*s*) := d*N* (*s*)*/*d*s. Adopt the c-parametrized formalism in (2) and fix every system parameter except c. If the open-loop* 𝒳 *satisfies the following conditions, then there exists a sufficiently small c*^∗^ *such that for every* 0 *< c* ≤ *c*^∗^ *the equilibrium point* Σ *of the closed-loop system (1) is locally asymptotically stable*.

*(C1) The given open-loop network* 𝒳 *is stable*

*(C2) σ*_11_*cρµη*_1_*N* (0) *>* 0

*(C3)* 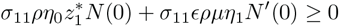

*Proof*. The controller subsystem involves an integrator, which is unstable, suggesting that the small-gain theorem may not be applicable to the closed-loop interconnection. We rely instead on an argument based on the root locus rules and perturbation of complex-valued functions to provide our proof. The characteristic polynomial of the corresponding linearized system can be written as 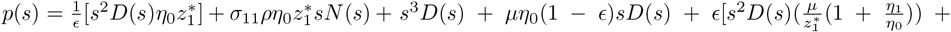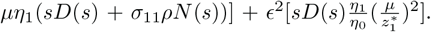. See Supplementary Material, Section S2 for detailed derivations. From the form of *p*(*s*), it can be seen that *n* + 2 of its roots approach to those of the dominant term *s*^2^*D*(*s*) as ϵ → 0. Thus, *n* of them are stable by (C1) as they are the roots of *D*(*s*). The root locus rules entail that the negative real axis is an asymptote, so there exists a stable root approaching −∞ in the limit ϵ → 0. This can be seen by recasting *p*(*s*) = 0 as 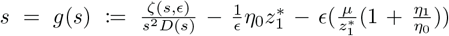,where ζ represents the residual terms. Since *P* (*s*) is proper, that is deg(*D*) ≥ deg(*N*), *s*^2^*D*(*s*) grows faster than ζ in the limit |*s*| → ∞ for any finite *c*. Hence, in this limit, the intersection of the functions *s* and *g*(*s*) tends to −∞ as ϵ tends to zero. Now consider an infinitesimally small disc *𝒟* = {*s* ∈ C : |*s*| ≤ *r*^∗^} with 0 *< r*^∗^ « 1 about the origin of the complex plane. Then, corresponding to every small *r*^∗^ there exists a sufficiently small *c*^∗^ such that for every ϵ ≤ ϵ ^∗^ two roots of *p*(*s*) lie on *𝒟*, while the remaining roots have all negative real parts. The proof will be complete if we find conditions that enforce these two roots both lie in the open left-hand half of *𝒟*.

We calculate the value of *p*(*s*) at *s* = 0 as *p*(*s*) _*s*=0_ = *σ*_11_*cρµη*_1_*N* (0). (A2) implies 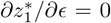.The derivative of *p*(*s*) w.r.t. *s*, which we know exists since it is a holomorphic function, evaluated at 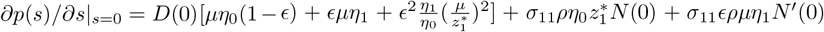 Note, 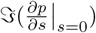 is equal to zero as *D, N*, and *N* ^′^ are polynomials of *s* with real-valued coefficients. Also, *D*(0) is real and positive. Moreover, the term 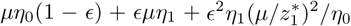 is always positive for ϵ *<* 1. It is easy to observe that *p*(0) is strictly positive for any ϵ *>* 0 if (C2) is met. Also, for any *c*, 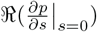is a positive number if (C3) is met. Equivalently, the phase of *p*(*s*) does not change local to the origin as *s* → 0 in any arbitrary direction. Therefore, as *r*^∗^ goes to zero, *p*(*s*) maps the points existing in the right-hand half of *𝒟* to the right of the vertical line ℜ(*p*(0)). These points, thus, cannot be roots of *p*(*s*), as ℜ(*p*(0)) *>* 0. Taken together, meeting (C1)-(C3) guarantees that the two roots in the interior of *𝒟* have always negative real parts as *r*^∗^ → 0.

### Remark 1.

*For small enough cs and when actuating positively (negatively) by* **Z**_1_, *or equivalently when σ*_11_ *is positive (negative), the process-dependent conditions (C2)-(C3) are always fulfilled regardless of N* ^t^(0), *provided N* (0) *>* 0 *(N* (0) *<* 0).

### Remark 2.

*Assume that (A1)-(A2) and (C1)-(C3) hold. As* ϵ → 0, *two closed-loop poles approach the origin (from the stable side) and the rest go to the open-loop poles and* −∞.

Considering positive actuation from Z_1_, the conditions that the open-loop has to satisfy are that it (i) needs to be stable and (ii) must have a positive DC gain (none or an even number of unstable zeros), Remark 1 says. Now, assume that the process parameters or dynamics (resp. the controller parameters, including the actuation gains) are uncertain and vary within a bounded 𝒞 ^1^ set Δ*P* (resp. Δ*C*). According to Theorem 1, there exists an ϵ « 1 which makes the entire uncertain closedloop system stable as long as Δ*P* and Δ*C* do not violate (A1)-(A2) and Δ*P* does not make the open-loop 𝒳 unstable or its DC gain negative. Of note, structurally stable open-loop circuits with positive DC gain fulfill these conditions by structure, regardless of their exact parameter values. These include uncertain activation cascades of any size depicted in Fig. 3A, which follow 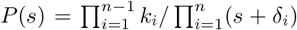.For any positive *k*_*i*_ and *δ*_*i*_, such open-loop circuits remain stable with a positive DC-gain given by 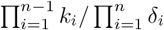.

Our numerical results provided in Fig. 4 support the findings above. Figs. 4A and C show that decreasing *c* improves the relative stability of the uncertain closed-loop circuit. Fig. 4A suggests that slightly decreasing *c* from 1 to ≈0.993 enables stable regulation over a substantially wider range of set-points.

**Fig. 4.**
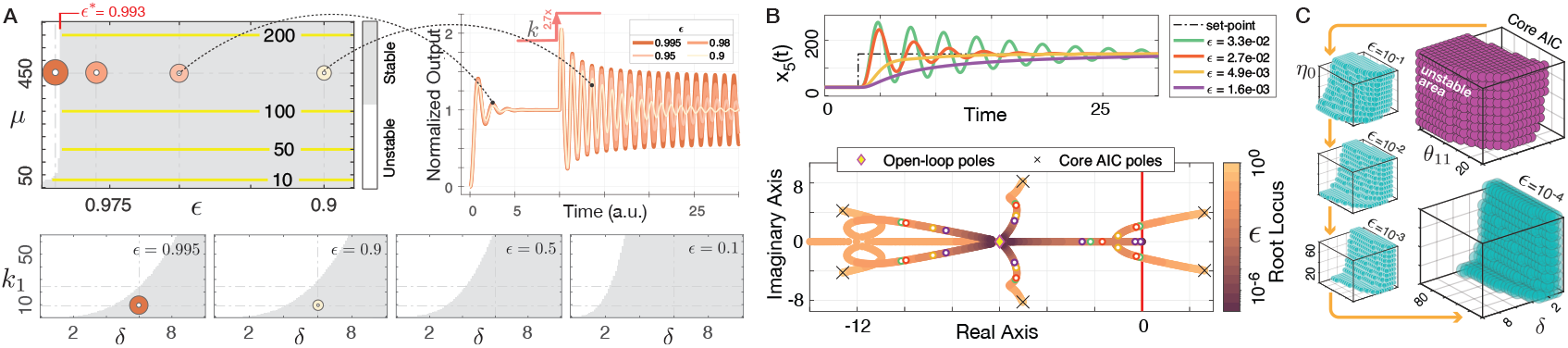
Cascaded antithetic integrators can be harnessed to improve stability robustness against uncertainties by adjusting the controller constant inflow production rates. We numerically evaluate the local stability of the closed-loop dynamic model from (1), where the actuation function Θ follows Θ = *θ*_11_*z*_1_. (A) Compared to the core AIC motif (when *ϵ* = 1), decreasing *ϵ* improves the closed-loop relative stability for variations in the set-point and process parametric uncertainties (*n* is set to two). Contour lines in yellow indicate the corresponding set-point. The 2D stability plots show the stability status over an uncertain 2D parametric region. (B) Root-locus plot of the closed-loop system with *n* = 5 for variations in *ϵ*. Corresponding set-point tracking responses for four sample *ϵ*s are shown in the top row. (C) The 3D stability plot for simultaneous bounded parametric uncertainties in both controller and process dynamics. The three parameters *δ, θ*_11_, and *η*_0_ are varied (*n* = 5). *δ* represents the constant degradation of every process species, meaning each *δ*_*i*_ = *δ*.

As verified by the stability indices in Fig. 3C, for varying number of process species, *n*, choosing *c* below ≈10^−4^ consistently stabilizes the entire system despite large parametric uncertainties. According to Theorem 1 and Remark 2, two closed-loop poles approach the origin as we improve the robustness by pushing *c* toward zero. This means that the system response becomes slower in the limit ϵ → 0, resulting in an overdamped response with no overshoot that is highly robust to uncertainties. This is consistent with the robustness-performance trade-off discussed earlier and is confirmed by results in Fig. 3 and Fig. 4B. We note that while generalizing our theorem to cases where (A2) is modified to allow actuations from **Z**_**2**_ and **Z**_**3**_ may follow similar steps and appear straightforward, it could lead to convoluted process-dependent conditions, making it difficult to draw general stability conclusions based solely on the characteristics of 𝒳.

## V. Biomolecular Realizations Using Genetic Parts

Here, using synthetic genetic parts we explore different realizations of our proposed control strategies considered in previous sections. We depict in Fig. 5 two of these realizations, one representing a PI and the other PID motif as discussed in Section III. They utilize inteins—short amino acid sequences capable of protein splicing [16]—to implement the sequestration mechanisms. Further circuits along with relevant details are provided as a note in Supplementary Material, Section S4.

**Fig. 5.**
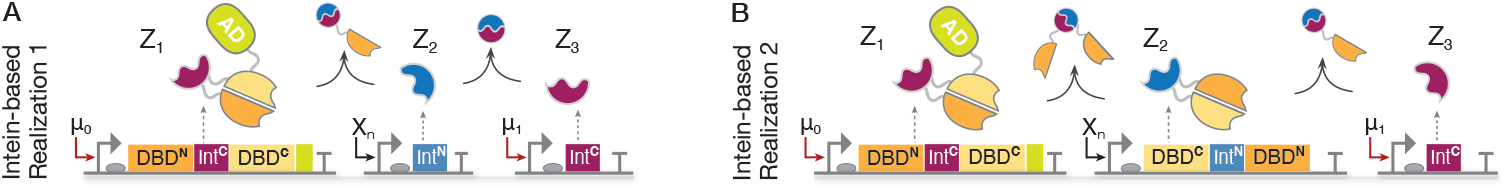
Biomolecular realizations of the minimal cascaded antithetic integrators through protein-protein sequestration reactions mediated by split inteins. We utilize an intein-mediated framework, introduced in [16] to realize the core AIC motif, to suggest cascading several annihilation modules. (A) represents a PI control structure in which only **Z**_**1**_ is transcriptionally active. By fusing more activation domains (AD) or changing the activation strength, one can tune its affinity for acting on a downstream gene. For instance, **Z**_**1**_ can serve as a transcription factor for the promoter of an upstream gene that regulates **X**_**1**_, establishing an intermediary positive actuation on the process. The circuit in (B) realizes a PID control structure by allowing **Z**_**2**_ to act as a repressor for **X**_**1**_.

## VI. Conclusion

Promising for synthetic biology, precise regulation and synthetic adaptation can be achieved using biomolecular integral controllers. In this letter, we considered cascaded AICs. Compared to the core AIC, our proposed integrators encode various PI and PID control structures, which we showed can be exploited to improve dynamic performance. Our numerical results verified this across a range of parameters (Fig. 2). Considering uncertainties in the nominal system, our mathematical analysis showed that our proposed control structure provides accessible means to improve robust stability in controlling a wide class of uncertain CRNs. This could be further leveraged to shape the governing robustness-performance trade-off (Fig. 3). Lastly, we discussed potential genetic implementations of our introduced motifs within synthetic circuits.

