## Supplementary Material for "Cascaded Antithetic Integral Feedback for Enhanced Stability and Performance"

### Outline

In this Supplementary Material for the manuscript "*Cascaded Antithetic Integral Feedback for Enhanced Stability and Performance*," we first discuss the dynamical model associated with the generalized cascaded antithetic integrators. Next, we provide a note on the linear perturbation analysis related to the minimal representation of these motifs. Finally, we include the parameter values used to obtain the numerical results and simulations presented in the main manuscript.

### S1. The Generalized Cascaded Antithetic Motif and its Associated Dynamical Model

According to the descriptions of the controller motif provided in the main text, we consider a generalized cascaded AIC motif up to the $q$-th order. We write down the following closed-loop model comprising $q \geq 3$ distinct controller species $\mathbf{Z_1}$ to $\mathbf{Z_q}$

$$\begin{cases} \dot{x} = \mathcal{F}(x) + \overbrace{e_1\Theta(z_1, \ldots, z_q, x_1)}^{\text{control input}}, & \text{(S1a)} \\ \dot{z}_1 = \mu_0 - \eta_0 z_1 z_2, & \text{(S1b)} \\ \dot{z}_2 = \rho x_n - \eta_0 z_1 z_2 - \eta_1 z_2 z_3, & \text{(S1c)} \\ \dot{z}_3 = \mu_1 - \eta_1 z_2 z_3 - \eta_2 z_3 z_4, & \text{(S1d)} \\ \vdots \quad \vdots \quad \vdots \quad \vdots & \\ \dot{z}_{q-1} = \mu_{q-3} - \eta_{q-3} z_{q-2} z_{q-1} - \eta_{q-2} z_{q-1} z_q, & \text{(S1e)} \\ \dot{z}_q = \mu_{q-2} - \eta_{q-2} z_{q-1} z_q, & \text{(S1f)} \end{cases}$$

where the state variable $x = [x_1, \cdots, x_n] \in \mathbb{R}^n_{\geq 0}$ represents the process species' concentrations, and the function $\mathcal{F}$ determines how dynamically they influence each other. The state variables $z_i$ denote the concentration of the controller species $\mathbf{Z_i}$ at time $t$. It is our assumption that the free actuation term $\Theta$ is properly engineered to ensure that the resulting closed-loop system admits a fixed point in the positive orthant $\mathbb{R}^{n+q}_{\geq 0}$ for the command set-point $(\mu_0 + \sum_{i=1}^{q-2}(-1)^{i-1}\mu_i)/\rho$. Let us, hence, assume that the process network and the dynamics of $\Theta$ determine a positive steady-state solution for $z_1^*$ standalone. Having a positive value for $z_1^*$, solving from (S1b) determines $z_2^* = \mu_0/\eta_0 z_1^*$. We use the superscript notation "$*$" to denote the corresponding steady-state solutions. Other controller species are not guaranteed to take positive values, yet. Indeed, how we choose the constant inflows $\mu_i$, even though it does not change the set-point, may still lead to incompatibility in admitting the corresponding positive equilibrium. Regarding this, a set of necessary conditions must be fulfilled for this admissibility, which we will be listing them in what follows.

These conditions arise solely from the controller dynamics and impose constraints on the constant inflow rates $\mu_i$. To list them, let us first define some new constants $\beta_i$ in terms of $\mu_i$ as follows

$$\beta_i \coloneqq \mu_i - \beta_{i+1} \qquad \text{for } 1 \leq i \leq q-3, \tag{S2}$$

$$\beta_{q-2} \coloneqq \mu_{q-2}. \tag{S3}$$

Note that each $\beta_i$ can be recursively calculated from the above formulae solely in terms of $\mu_i$s. In particular, $\beta_1 \coloneqq \sum_{i=1}^{i=q-2}(-1)^{i-1}\mu_i$. Solving (S1b)-(S1f) for the equilibria results in

$$\eta_i z_{i+1}^* z_{i+2}^* = \beta_i, \tag{S4}$$

for $i \in \{q-2, q-3, \ldots, 2, 1\}$. Taken together, one can observe that all the necessary conditions for positive equilibria imposed by the controller are satisfied if all the constants $\beta_i$ are strictly positive for $i \in \{1, \ldots, q-2\}$. This entails that, for every $i \in \{3, \cdots, q-1\}$, each $\mathbf{Z_i}$ must be constitutively produced at a rate higher than its adjacent $\mathbf{Z_{i+1}}$ for the system to admit the set-point $(\mu_0 + \beta_1)/\rho$. One can follow this from the constraint $\mu_i - \mu_{i+1} + \beta_{i+2} > 0$ that has to be held for $i \in \{1, \cdots, q-4\}$. For the minimal case where $q = 3$, there is no necessary condition for the admissibility of a positive equilibrium imposed from the controller side. That is, for every positive arrangement of $\mu_0$ and $\mu_1$, $z_1^*$, $z_2^*$, and $z_3^*$ always admit a positive steady-state solution. Of course, there may always exist additional conditions imposed by the process dynamics that need to be satisfied to make the set-point admissible. These conditions should be checked for a specific process.

### S2. Linear Perturbation Analysis

In this section, we are concerned with the following closed-loop dynamic model

$$\begin{cases} \dot{x} = \mathcal{F}(x) \overbrace{+ e_1 \Theta(z_1, z_2, z_3),}^{\text{free actuation terms}} & \text{(S5a)} \\ \dot{z}_1 = \epsilon\mu(t) - \eta_0 z_1 z_2, & \text{(S5b)} \\ \dot{z}_2 = \rho x_n - \eta_0 z_1 z_2 - \eta_1 z_2 z_3, & \text{(S5c)} \\ \dot{z}_3 = (1-\epsilon)\mu(t) - \eta_1 z_2 z_3, & \text{(S5d)} \end{cases}$$

where $\mu(t) \in \mathbb{R}_{\geq 0}$ now represents the reference input signal. By performing a linear small perturbation analysis on this closed-loop dynamic model, we report the underlying characteristic polynomial and associated PI/PID gains. Note that, when $\epsilon = 1$, the controller's response reduces to that of the corresponding core AIC motif (provided the initial condition for $z_3$ is zero).

#### A. Linearized Dynamics of the Closed-loop System and Its Characteristic Polynomial

In this subsection, we aim to obtain the characteristic polynomial of the system (S5) with dynamics linearized about the fixed point $\Sigma = [x^*, z_1^*, z_2^*, z_3^*] \in \mathbb{R}_{\geq 0}^{n+3}$, where $x^* \coloneqq [x_1^*, \cdots, x_n^*]$. We denote this characteristic polynomial as $p(s)$, where $s \in \mathbb{C}$. It is our assumption that such a non-negative fixed point $\Sigma$ exists. For a constant $\mu(t) = \mu$, (S5) follows that

$$x_n^* = \mu/\rho, \tag{S6}$$

$$z_2^* = \epsilon\mu/\eta_0 z_1^*, \tag{S7}$$

$$z_3^* = (1/\epsilon - 1)\eta_0 z_1^*/\eta_1. \tag{S8}$$

Note that $z_3^* \to \infty$ as $\epsilon \to 0$. We will perform a linear perturbation analysis. To begin with, we introduce the following new variables for $i \in \{1, \ldots, n\}$ and $j \in \{1, 2, 3\}$

$$\tilde{x}_i(t) = x_i(t) - x_i^*,$$

$$\tilde{z}_j(t) = z_j(t) - z_j^*,$$

$$\tilde{\mu}(t) = \mu(t) - \mu,$$

where the actual input $\mu(t)$, representing the set-point to be tracked, is allowed to have small variations about the fixed value $\mu$. We use non-bold capital letters to signify the corresponding Laplace transform of a time-domain signal, denoted by lowercase letters.

Let us write down the following relationships for the linearized dynamics of the closed-loop model (S5)

$$\tilde{X}_n(s) = P(s)\tilde{\Theta}(s), \tag{S9}$$

$$\tilde{\Theta}(s) = \overbrace{\begin{bmatrix}\sigma_{11} & \sigma_{21} & \sigma_{31}\end{bmatrix}}^{\sigma} \begin{bmatrix}\tilde{Z}_1(s) \\ \tilde{Z}_2(s) \\ \tilde{Z}_3(s)\end{bmatrix}, \tag{S10}$$

$$\begin{bmatrix}\tilde{Z}_1(s) \\ \tilde{Z}_2(s) \\ \tilde{Z}_3(s)\end{bmatrix} = \underbrace{\begin{bmatrix}C_{11}(s) \\ C_{12}(s) \\ C_{13}(s)\end{bmatrix}}_{C_n(s)} \tilde{X}_n(s) + \underbrace{\begin{bmatrix}C_{21}(s) \\ C_{22}(s) \\ C_{23}(s)\end{bmatrix}}_{C_\mu(s)} \tilde{\mu}(s). \tag{S11}$$

$\tilde{\Theta}(s)$ is the Laplace transform of the small-signal linearization of the control input signal applied to the plant $\boldsymbol{\mathcal{X}}$. These equalities result in the following relation

$$\frac{\tilde{X}_n(s)}{\tilde{\mu}(s)} = \frac{P\sigma C_\mu}{1 - P\sigma C_n}. \tag{S12}$$

We provide the associated block diagram schematic of this model in Fig. S1A. We treat the process-related transfer function $P(s)$ as an unknown but rational transfer function, following the formula $P(s) := N(s)/D(s)$, where $N$ and $D$ are polynomials in $s$. This transfer function follows $P(s) = e_n^{\mathrm{T}} \left(sI - A_{\mathrm{p}}\right)^{-1} e_1$, where $A_{\mathrm{p}} := \partial\mathcal{F}|_\Sigma$ and $I$ is the identity matrix of appropriate size. Note that $D(s)$ represents the open-loop characteristic polynomial. The transfer function $P(s)$ is expected to be proper which remains unchanged as long as the process parameters and structure do not change.

The potentially $\epsilon$-dependent terms $\sigma_{i1}$ are constants, which could be positive, negative, or zero, depending on the biomolecular reactions governing the actuation from the controller side. They represent the linearized effect of the actuation terms from the species $\mathbf{Z_i}$ to $\mathbf{X_1}$, so that

$$\sigma_{i1} := \partial\Theta(z_1, z_2, z_3, x_1)/\partial z_i\big|_{(z_1,z_2,z_3,x_1)=q}.$$

Thus, if the linearized effect of the actuation from $\mathbf{Z_i}$ to the process input reflected through the function $\Theta$ is negative, noninfluential, or positive, we expect the term $\sigma_{i1}$ to be negative definite, zero, or positive definite, respectively.

Now, we derive the expressions for the controller transfer functions $C_{ij}(s)$, given the dynamics on the controller side. To that end, we need to compute each $C_{ij}$ separately. For $C_{11}(s)$, we find

$$\begin{aligned} C_{11}(s) &= \frac{\tilde{Z}_1(s)}{\tilde{X}_n(s)}\bigg|_{\tilde{\mu}(s)=0} \\ &= \begin{bmatrix} 1 & 0 & 0 \end{bmatrix} (sI - A)^{-1} B^{\mathrm{T}}, \end{aligned} \tag{S13}$$

with $I$ being the identity matrix of an appropriate size and $A$ defined as being the Jacobian matrix of the controller dynamics evaluated at $q$, which follows

$$A = \begin{bmatrix} -\eta_0 z_2^* & -\eta_0 z_1^* & 0 \\ -\eta_0 z_2^* & -\eta_0 z_1^* - \eta_1 z_3^* & -\eta_1 z_2^* \\ 0 & -\eta_1 z_3^* & -\eta_1 z_2^* \end{bmatrix}. \tag{S14}$$

The vector $B \in \mathbb{R}^3$ is given by $B := \begin{bmatrix} 0 & \rho & 0 \end{bmatrix}$. By slight abuse of notation, we define

$$\alpha_0 := \eta_0 z_2^*, \quad \alpha_1 := \eta_1 z_3^*, \tag{S15}$$

$$\beta_0 := \eta_0 \mu_0, \quad \beta_1 := \eta_1 \mu_1, \tag{S16}$$

where $\mu_0 := \epsilon\mu$ and $\mu_1 := (1-\epsilon)\mu$. From the above definitions, one can note the equalities $\alpha_0 = \mu_0/z_1^*$, $\alpha_1 = \mu_1/z_2^* \equiv (1/\epsilon - 1)\eta_0 z_1^*$, $\beta_0/\alpha_0 = \eta_0 z_1^*$, $\beta_1/\alpha_1 = \eta_1 z_2^* = \eta_1\mu_0/(\eta_0 z_1^*)$, $\beta_0\beta_1/\alpha_0\alpha_1 = \eta_1\mu_0$, and $\alpha_0\beta_1/\alpha_1 = \eta_1\mu_0^2/\eta_0 {z_1^*}^2$. We rewrite $A$ as

$$A = \begin{bmatrix} -\alpha_0 & -\beta_0/\alpha_0 & 0 \\ -\alpha_0 & -\beta_0/\alpha_0 - \alpha_1 & -\beta_1/\alpha_1 \\ 0 & -\alpha_1 & -\beta_1/\alpha_1 \end{bmatrix}$$

and $C_{11}(s)$ as

$$C_{11}(s) = -\frac{\rho\beta_0}{\alpha_0}\frac{s + \beta_1/\alpha_1}{\Delta(s)},$$

where

$$\Delta(s) := s(s^2 + (\alpha_0 + \alpha_1 + \frac{\beta_0}{\alpha_0} + \frac{\beta_1}{\alpha_1})s + \alpha_0\alpha_1 + \alpha_0\frac{\beta_1}{\alpha_1} + \frac{\beta_0\beta_1}{\alpha_0\alpha_1}).$$

Note the factor of $1/s$ in $C_{11}(s)$ which is an indicator of the integrator.

Similar to the calculation of $C_{11}(s)$ from Section S2-A, we find the following for the other $C_{ij}(s)$

$$
\begin{aligned}
C_{12}(s) &= \frac{\tilde{Z}_2(s)}{\tilde{X}_n(s)}\Big|_{\tilde{\mu}(s)=0} = \begin{bmatrix} 0 & 1 & 0 \end{bmatrix} (sI - A)^{-1} \begin{bmatrix} 0 \\ \rho \\ 0 \end{bmatrix} \\
&= \rho \frac{(s+\alpha_0)(s+\beta_1/\alpha_1)}{\Delta(s)}, \\
C_{13}(s) &= \frac{\tilde{Z}_3(s)}{\tilde{X}_n(s)}\Big|_{\tilde{\mu}(s)=0} = \begin{bmatrix} 0 & 0 & 1 \end{bmatrix} (sI - A)^{-1} \begin{bmatrix} 0 \\ \rho \\ 0 \end{bmatrix} \\
&= -\rho \frac{\alpha_1(s+\alpha_0)}{\Delta(s)}, \\
C_{21}(s) &= \frac{\tilde{Z}_1(s)}{\tilde{\mu}(s)}\Big|_{\tilde{X}_n(s)=0} = \begin{bmatrix} 1 & 0 & 0 \end{bmatrix} (sI - A)^{-1} \begin{bmatrix} \epsilon \\ 0 \\ 1-\epsilon \end{bmatrix} \\
&= \frac{\epsilon(s+\frac{\beta_0}{\alpha_0})(s+\frac{\beta_1}{\alpha_1}) + \epsilon\alpha_1 s + \frac{\beta_0\beta_1}{\alpha_0\alpha_1}(1-\epsilon)}{\Delta(s)}, \\
C_{22}(s) &= \frac{\tilde{Z}_2(s)}{\tilde{\mu}(s)}\Big|_{\tilde{X}_n(s)=0} = \begin{bmatrix} 0 & 1 & 0 \end{bmatrix} (sI - A)^{-1} \begin{bmatrix} \epsilon \\ 0 \\ 1-\epsilon \end{bmatrix} \\
&= -\frac{\epsilon\alpha_0(s+\frac{\beta_1}{\alpha_1}) + \frac{\beta_1}{\alpha_1}(1-\epsilon)(s+\alpha_0)}{\Delta(s)}, \\
C_{23}(s) &= \frac{\tilde{Z}_3(s)}{\tilde{\mu}(s)}\Big|_{\tilde{X}_n(s)=0} = \begin{bmatrix} 0 & 0 & 1 \end{bmatrix} (sI - A)^{-1} \begin{bmatrix} \epsilon \\ 0 \\ 1-\epsilon \end{bmatrix} \\
&= \frac{\epsilon\alpha_0\alpha_1 + (1-\epsilon)\left[(s+\alpha_0)(s+\alpha_1) + \frac{\beta_0}{\alpha_0}s\right]}{\Delta(s)}.
\end{aligned}
$$

Recall that $\beta_0/\alpha_0 = \eta_0 z_1^*$ and $\beta_1/\alpha_1 = \epsilon\eta_1\mu/\eta_0 z_1^* \equiv \eta_1\alpha_0/\eta_0$.

Now, we are in position to write out the characteristic polynomial of the concerned closed-loop system. We shall call it by $p(s)$. From (S12), we know that the closed-loop follows

$$
\frac{\tilde{X}_n(s)}{\tilde{\mu}(s)} = \frac{\sum_{i=1}^{3} \sigma_{i1} N_{C_{2i}}(s) N(s)}{D(s)\Delta(s) - \sum_{i=1}^{3} \sigma_{i1} N_{C_{1i}}(s) N(s)}, \tag{S17}
$$

where $N_{C_{ij}}$ represents the numerator of the transfer function $C_{ij}$. Let us calculate this characteristic polynomial under the assumption that the controller acts on the plant only through $\mathbf{Z_1}$. This equivalently means to set $\sigma_{21} = 0$ and $\sigma_{31} = 0$, which reads $\tilde{\Theta}(s) = \sigma_{11}\tilde{Z}_1(s)$. By this assumption $z_1^*$ and $\sigma_{11}$ will become independent of $\epsilon$. Following this, $p(s) = D(s)\Delta(s) - \sigma_{11}N(s)N_{C_{11}}(s)$. Substituting from the above derivations and after some rearrangements, we can write out

$$\begin{aligned}
p(s) = & \frac{1}{\epsilon}[s^2 D(s)\eta_0 z_1^*] + \sigma_{11}\rho\eta_0 z_1^* sN(s) \\
& + s^3 D(s) + \mu\eta_0(1-\epsilon)sD(s) \\
& + \epsilon[s^2 D(s)(\frac{\mu}{z_1^*}(1+\frac{\eta_1}{\eta_0})) + \mu\eta_1(sD(s) + \sigma_{11}\rho N(s))] \\
& + \epsilon^2[sD(s)\frac{\eta_1}{\eta_0}(\frac{\mu}{z_1^*})^2].
\end{aligned} \tag{S18}$$

*B. Linearized Dynamics of the Standalone Controller Follow Different PI and PID Control Structures Depending on the Arrangement of Actuation Channels*

We now write out the control input $\tilde{\Theta}(s) \coloneqq \sum_{i=1}^{3} \sigma_{i1}\tilde{Z}_i(s)$ by assuming that all the actuation signals are applied to the same input species $\mathbf{X_1}$. We know that $\tilde{Z}_i(s) = C_{1i}\tilde{X}_n(s) + C_{2i}\tilde{\mu}(s)$. Define the tracking error term $\tilde{E}(s)$ as $\tilde{E}(s) \coloneqq \tilde{\mu}(s) - \rho\tilde{X}_n(s)$. Moreover, define the cut-off frequencies $\omega_1$ and $\omega_2$ as

$$\omega_{1,2} \coloneqq \frac{1}{2} \times \left( (\alpha_0 + \alpha_1 + \frac{\beta_0}{\alpha_0} + \frac{\beta_1}{\alpha_1}) \pm \sqrt{(\alpha_0 + \alpha_1 + \frac{\beta_0}{\alpha_0} + \frac{\beta_1}{\alpha_1})^2 - 4(\alpha_0\alpha_1 + \alpha_0\frac{\beta_1}{\alpha_1} + \frac{\beta_0\beta_1}{\alpha_0\alpha_1})} \right). \tag{S19}$$

By combining the above derivations, one can see that

$$\begin{aligned}
\tilde{\Theta}(s) = & \frac{\sigma_{11}}{s(s+\omega_1)(s+\omega_2)} \left[ \left( \epsilon(s + \frac{\beta_0}{\alpha_0})(s + \frac{\beta_1}{\alpha_1}) + \epsilon\alpha_1 s + \frac{\beta_0\beta_1}{\alpha_0\alpha_1}(1-\epsilon) \right) \tilde{\mu}(s) - \frac{\rho\beta_0}{\alpha_0}(s + \frac{\beta_1}{\alpha_1})\tilde{X}_n(s) \right] \\
& + \frac{-\sigma_{21}}{s(s+\omega_1)(s+\omega_2)} \left[ \left( \epsilon\alpha_0(s + \frac{\beta_1}{\alpha_1}) + \frac{\beta_1}{\alpha_1}(1-\epsilon)(s+\alpha_0) \right) \tilde{\mu}(s) - \rho(s+\alpha_0)(s + \frac{\beta_1}{\alpha_1})\tilde{X}_n(s) \right] \\
& + \frac{\sigma_{31}}{s(s+\omega_1)(s+\omega_2)} \left[ \left( \epsilon\alpha_0\alpha_1 + (1-\epsilon)\left( (s+\alpha_0)(s+\alpha_1) + \frac{\beta_0}{\alpha_0}s \right) \right) \tilde{\mu}(s) - \rho\alpha_1(s+\alpha_0)\tilde{X}_n(s) \right].
\end{aligned}$$

One can recast it as

$$\tilde{\Theta}(s) = \frac{\omega_1}{s+\omega_1}\frac{\omega_2}{s+\omega_2}\left( \frac{K_I}{s}\tilde{E}(s) + (K_{F_0} + K_{F_D}s)\tilde{\mu}(s) - (K_P + K_D s)\tilde{X}_n(s) \right),$$

where

$$\begin{aligned}
K_I &= (\sigma_{11}\frac{\beta_0\beta_1}{\alpha_0\alpha_1} - \sigma_{21}\frac{\alpha_0\beta_1}{\alpha_1} + \sigma_{31}\alpha_1\alpha_0)/\omega_1\omega_2 \\
K_{F_0} &= \left( \sigma_{11}\epsilon(\alpha_1 + \frac{\beta_0}{\alpha_0} + \frac{\beta_1}{\alpha_1}) - \sigma_{21}\left( \epsilon\alpha_0 + \frac{\beta_1}{\alpha_1}(1-\epsilon) \right) + \sigma_{31}(1-\epsilon)(\alpha_0 + \alpha_1 + \frac{\beta_0}{\alpha_0}) \right)/\omega_1\omega_2 \\
K_{F_D} &= (\sigma_{11}\epsilon + \sigma_{31}(1-\epsilon))/\omega_1\omega_2 \\
K_P &= \rho\left(\sigma_{11}\beta_0/\alpha_0 - \sigma_{21}(\alpha_0 + \beta_1/\alpha_1) + \sigma_{31}\alpha_1\right)/\omega_1\omega_2 \\
K_D &= -\sigma_{21}\rho/\omega_1\omega_2.
\end{aligned}$$

By definition, $\alpha_i$ and $\beta_i$ are strictly positive scalars for every $0 < \epsilon < 1$ (of course, under the assumption of set-point admissibility). Recall that the constants $\sigma_{i1}$s are sign-indefinite. Note that $\omega_1\omega_2$ equals to the value $\alpha_0\alpha_1 + \alpha_0\beta_1/\alpha_1 + \beta_0\beta_1/\alpha_0\alpha_1$. The P and D components act on the output feedback. The block diagram schematic of the input-output signals corresponding to the controller is depicted in Fig. S1B. As can be easily seen from the above gains, for non-negative, non-positive, and non-negative actuation from $\mathbf{Z_1}$, $\mathbf{Z_2}$, and $\mathbf{Z_3}$, respectively, we expect having all gains positive, including $K_{F_0}$ and $K_{F_D}$.

Different arrangements of the actuation channel might alter the control structure from a filtered PI to PID. We categorize seven different rearrangements of such, reported in Fig. S2. These controllers do not introduce additional closed-loop unstable zeros. To see this, let us write down the formula for the signal which maps the reference input to $\tilde{X}_n(s)$. This way, we arrive at this expression

$$\tilde{\Theta}(s) = \frac{N_\mu}{D_\mu}\tilde{\mu}(s) + \frac{N_x}{D_x}\tilde{X}_n(s),$$

wherein $N_\mu = \omega_1\omega_2(K_I + K_{F_0}s + K_{F_D}s^2)$, $N_x = -\omega_1\omega_2(\rho K_I + K_P s + K_D s^2)$, and $D_\mu = D_x = s(s+\omega_1)(s+\omega_2) =: D_c$. Next, from (S9) it can be found that

$$\frac{\tilde{X}_n(s)}{\tilde{\mu}(s)} = \frac{N(s)N_\mu(s)}{D(s)D_c(s) - N(s)N_x(s)}.$$

As expected, the feedforward zeros have remained intact in the closed-loop transfer function. Thus, if the roots of the term $N_\mu(s)$ are always stable, then we expect that the linearized dynamics of the controller standalone cannot lead to a non-minimum phase behavior in the closed-loop system (given a minimum-phase process). We can claim so if $K_I$, $K_{F_0}$, and $K_{F_D}$ are all positive.

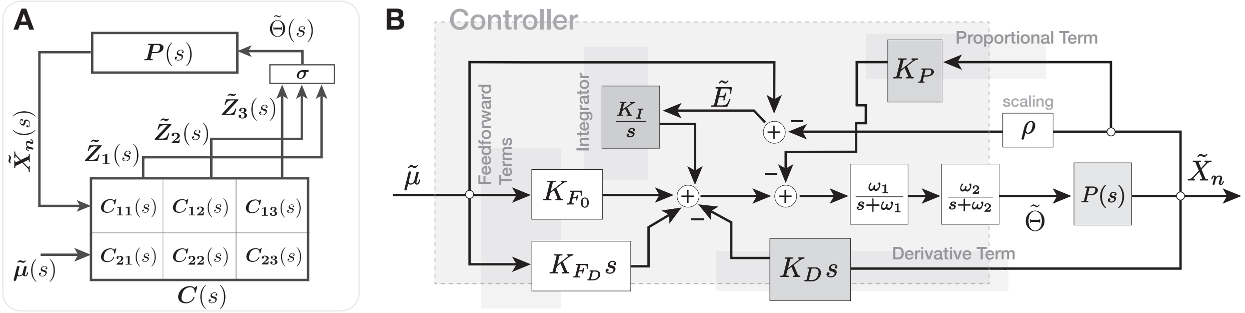


Fig. S1. (A) Schematic block diagram describing the linearized dynamics of the closed-loop system (S5). (B) The block diagram of the cascaded AIC controller standalone, $C(s)$. In line with the discussion in Section S2-B, the linear small perturbation model of the controller encodes different PI and PID control structures upon specific arrangements of the actuation terms through $\Theta$.

### S3. Numerical Values for the Parameters

Here we report the numerical values used to obtain our simulation results presented as Fig. 2 to Fig. 4. The initial conditions were set to zero. The regulated process follows the formulation of an $n$-species linear activation cascade ($n \geq 2$), where each species $\mathbf{X_i}$ positively influences the production of the adjacent species $\mathbf{X_{i+1}}$. Define the positive parameters $k_i$ and $\delta_i$ to be the process production and degradation rate constants, respectively. The

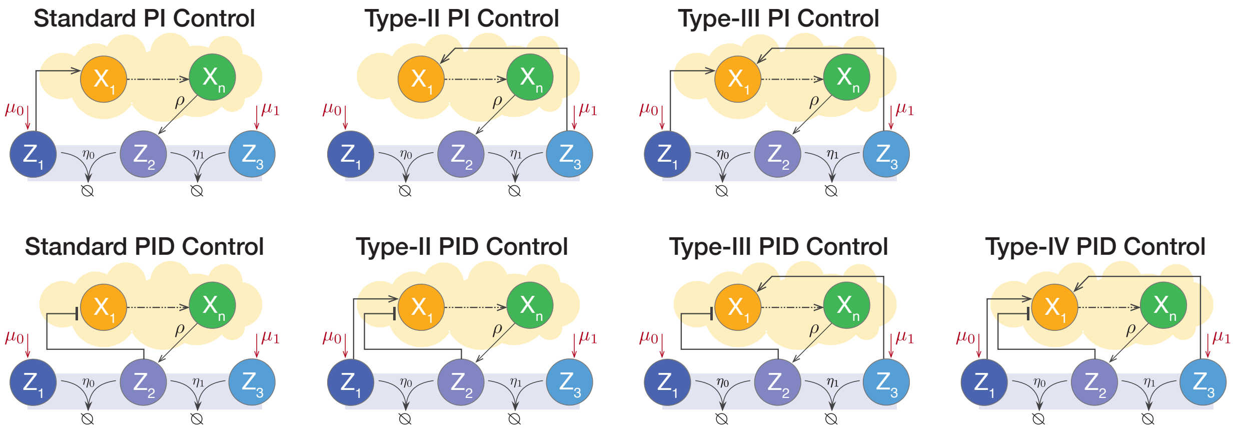


Fig. S2. Different arrangements of actuation channels might lead to different control structures (filtered PI or PID) in the minimal cascaded AIC. We illustrate seven such arrangements, three of which correspond to a filtered PI and the rest to a filtered PID formalism. The corresponding gains for these seven motifs are positive by structure, and they do not introduce nonminimum-phase closed-loop zeros additional to those that the open-loop network does.

controller network follows the formulation of the minimal cascaded AIC. The dynamical formulation of the closed-loop circuit is as follows

$$\begin{cases} \dot{x} = Ax + e_1\Theta(z_1, z_2, z_3), & \text{(S20a)} \\ \dot{z}_1 = \epsilon\mu - \eta_0 z_1 z_2, & \text{(S20b)} \\ \dot{z}_2 = \rho x_n - \eta_0 z_1 z_2 - \eta_1 z_2 z_3, & \text{(S20c)} \\ \dot{z}_3 = (1-\epsilon)\mu - \eta_1 z_2 z_3, & \text{(S20d)} \end{cases}$$

where $A := [a_{ij}] \in \mathbb{R}^{n\times n}$ is a lower-triangular matrix with negative diagonals. Specifically, $a_{ii}$ are set to $-\delta_i$ for every $i$, and for $i \in \{1, \cdots, n-1\}$, each $a_{(i+1)i}$ is given by $a_{(i+1)i} := k_i$. The rest of the elements of $A$ are zero. The normalization factor for the output of interest, used in certain plots to obtain the normalized concentrations, was set to the set-point value, $\mu/\rho$. The computation of the overshoot values were also based on the normalized outputs. For example, a computed value $Y_{\text{ovr}} = 0.4$ corresponds to 40% overshoot from the baseline level $\mu/\rho$.

**Fig. 2:** The nominal parameters used were: $n = 2$, $\mu = 450$, $\eta_0 = 60$, $\eta_1 = 40$, $\rho = 3$, $\delta_1 = \delta_2 = 6$, and $k_1 = 9 \times 2.7$. Given that $\Theta := \theta_{31}z_3 + \theta_{11}z_1/(1 + \theta_{21}z_2)$, solving for the steady-state solution of $z_1$ yields the unique solution

$$z_1^* = \frac{-(\theta_{21}\theta_{31}\frac{\mu}{2\eta_1}(1-\epsilon) - \phi/2) + \sqrt{(\theta_{21}\theta_{31}\frac{\mu}{2\eta_1}(1-\epsilon) - \phi/2)^2 + \frac{\theta_{21}}{\eta_0}\epsilon\mu\phi(\theta_{11} + \theta_{31}\frac{\eta_0}{\eta_1}(1/\epsilon - 1))}}{\theta_{11} + \theta_{31}\eta_0(1/\epsilon - 1)/\eta_1},$$

where $\phi$ is defined as the supporting input the process needs to admit the set-point. $\phi$ depends on the process structure, parameters, and the set-point value. In the examples considered in this figure, $\phi$ is given by $\phi = \delta_1\delta_2\mu/\rho k_1$.

**Fig. 3:** The nominal parameters used were: $\mu = 450$, $\eta_0 = 60$, $\eta_1 = 40$, and $\rho = 3$. In these simulations, the parameter $n$, representing the number of process species, varies. We set $\delta_1$ and $\delta_2$ to 6, while the rest $\delta_i$ were set to unity. For the $k_i$s, all were set to unity except $k_1 = 9$.

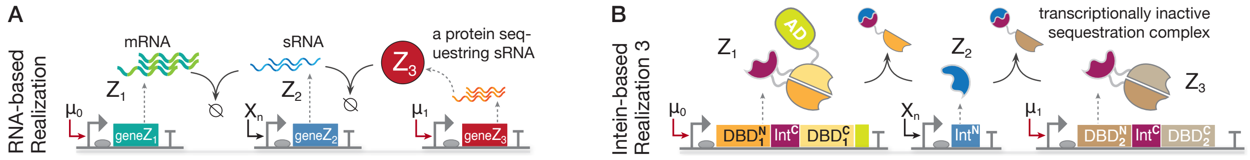


Fig. S3. Continuation of the biomolecular realizations of the minimal cascaded antithetic integrators through synthetic genes. (A) A specific mRNA may be sequestered by a cognate sRNA, which is simultaneously being sequestered by an RNA-binding protein. The resulting protein from the translation of this mRNA may act as a transcriptional activator or repressor for other genes, establishing the actuation in the same way as in Fig. 5A. (B) enables the realization of the type-III PI mechanism, now with $\mathbf{Z_3}$ being a transcriptionally active complex. Note that the species $\mathbf{Z_3}$ in (B) can act directly as a repressor for $\mathbf{X_1}$'s promoter or may establish an intermediary positive interaction with it, instead. As such, it may be engineered to repress an intermediate gene that is a repressor for $\mathbf{X_1}$.

**Fig. 4:** For the panel (A), the nominal parameters used were: $n = 2$, $\mu = 450$, $\eta_0 = 60$, $\eta_1 = 40$, $\rho = 3$, $\delta_1 = \delta_2 =: \delta = 6$, $k_1 = 9$, and $\theta_{11} = 6$. For the panels (B) and (C) they were: $n = 5$, $\eta_0 = 60$, $\eta_1 = 40$, $\rho = 3$, $\delta_i =: \delta = 6$, $k_1 = 9$, $k_2 = 6$, $k_3 = 9$, $k_4 = 6$, and $\theta_{11} = 90$. In the set-point tracking responses shown in (B), the top part, the set-point changes from 30 to 150 by changing $\mu$ from 90 to 450. The trajectories were already settled before the set-point changes.

### S4. Supplementary Notes and Simulation Results

#### A. A Complementary Note on Biomolecular Implementation and Natural Occurrence of the Proposed Designs

Reported in Table S1, one can find examples of cascaded sequestration in different bacteria based on the existing reports in the literature. In the main text, Section V, we explored two realizations of our proposed cascaded antithetic-based control circuits. Here, we provide two more example realizations. The first relies on non-coding small RNAs and a protein engineered to sequester them, complementing the cascade. This is depicted in Fig. S3A. The minimal cascaded AIC motif, which actuates only through $\mathbf{Z_1}$ and follows the standard PI structure, can be realized by this circuit in Fig. S3A. The type-III PI mechanisms, on the other hand, can be realized by the intein-mediated realization provided in Fig. S3B.

Upon proper protein folding, the split intein pairs dimerize and undergo protein *trans* splicing, which shuffles the sequences linked to them. This process covalently links the peptide upstream of $\text{Int}^{\text{N}}$ to the peptide downstream of $\text{Int}^{\text{C}}$, with the remaining sequences remaining as a separate heterodimer. This mode of action enables irreversible sequestration between the two protein complexes to which the intein pairs are fused, resulting in functional/inactive sequestration complexes. For more details, see [1], [2] and the references therein. In intein-mediated realizations, we have intentionally fused the split DNA binding domains (DBDs) in locations such that the resulting sequestration complexes are transcriptionally inactive. Note that the species with two complementary split DBDs fused to them can act as a repressor for a downstream gene if not attached to an activation domain (AD) or as an activator if attached to an AD. We have used this flexibility in the provided circuits to wire the actuation channels, thereby spanning the design to different PI and PID mechanisms discussed in Section III of the main text and Section S2 above.

TABLE S1

CASCADED FORMS OF SEQUESTRATION IN BACTERIAL PATHOGENS

| Mappings to the Motif in Fig. 1 | | | Regulatory Function in[d] | Bacterial Strain | Ref. |
|---|---|---|---|---|---|
| $\mathbf{Z_3}$[a] | $\mathbf{Z_2}$[b] | $\mathbf{Z_1}$[c] | | | |
| ExsC | ExsD | ExsA | T3SS | *P. aeruginosa* | [3], [4] |
| RsfA/B | UsfX | SigF | SACMM | *M. tuberculosis* | [5] |
| HrpG | HrpV | HrpR-HrpS | T3SS | *P. syringae* | [6] |
| InvE | SipC | InvF-SicA | T3SS | *S. typhimurium* | [7] |

[a] ExsC, RsfA/B, HrpG, and InvE: anti-anti-activators; ExsC has a secondary function as a chaperone; InvE is an invasion protein in Salmonella.

[b] ExsD, UsfX, HrpV, and SipC: anti-activators.

[c] ExsA: AraC-type transcription factor; SigF: alternate sigma factor; HrpR, HrpS, and InvF: transcriptional activators; SicA: type-III secretion chaperone.

[d] T3SS: Type-III secretion system; SACMM: Stress adaptation and cell membrane modification.

### B. Supplementary Simulation Results

We provide in Fig. S4 numerical results for six of the seven motifs illustrated in Fig. S2. The nominal parameters and process network are as in Fig. 2 of the main text, with the only change being how the controller acts on the plant. The selected actuation function for all simulations in Fig. S4 is as follows: $\Theta = \theta_{11}z_1 - \theta_{21}z_2x_1 + \theta_{31}z_3$, implying the inhibitory reaction between $\mathbf{Z_2}$ and $\mathbf{X_1}$ is of active degradation type. Given the supporting input $\phi$, the steady-state solution of $z_1$ follows $z_1^* = \left(\phi/2 + \sqrt{(\phi/2)^2 + \frac{\theta_{21}}{\eta_0}\epsilon\mu\phi(\theta_{11} + \theta_{31}\frac{\eta_0}{\eta_1}(1/\epsilon - 1))/\delta_1}\right)/(\theta_{11} + \theta_{31}\eta_0(1/\epsilon - 1)/\eta_1)$. We set $\theta_{11} = 6$, $\theta_{21} = 50$, and $\theta_{31} = 1$ to obtain the simulations with initial conditions set to zero.

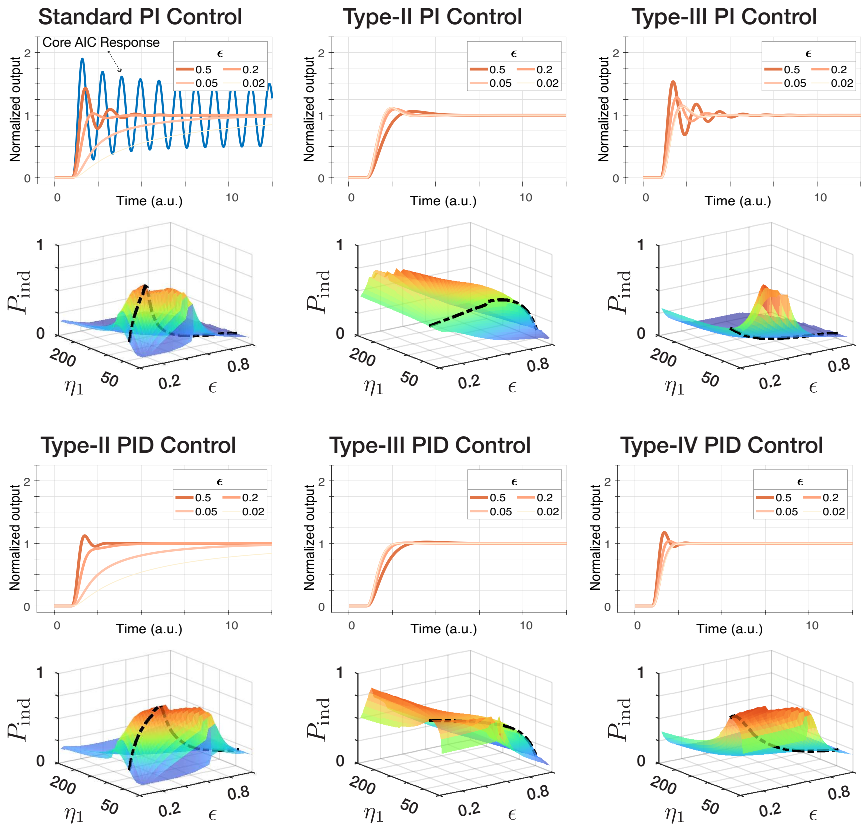


Fig. S4. Time response and performance index calculations for six out of the seven motifs depicted in Fig. S2. The regulated process is a two-species activation cascade (representing a simplified gene expression model). The pointwise performance indices are computed over a range of variations in $\epsilon$ and $\eta_1$, with $\epsilon$ ranging from 0 to 1 and $\eta_1$ from 1 to 250. Overall, PID controllers show better performance than PI motifs, as indicated by more orange and red regions across a wider parameter range.
